# Overcoming the design, build, test (DBT) bottleneck for synthesis of nonrepetitive protein-RNA binding cassettes for RNA applications

**DOI:** 10.1101/2019.12.24.886168

**Authors:** Noa Katz, Eitamar Tripto, Sarah Goldberg, Orna Atar, Zohar Yakhini, Yaron Orenstein, Roee Amit

**Affiliations:** Department of Biotechnology and Food Engineering, Technion - Israel Institute of Technology, Haifa 3200003, Israel; Department of Biomedical Engineering, Ben-Gurion University of the Negev, Beer-Sheva 8410501, Israel; Department of Computer Science, Technion - Israel Institute of Technology, Haifa 3200003, Israel; School of Computer Science, Interdisciplinary Center, Herzliya 46150, Israel; School of Electrical and Computer Engineering, Ben-Gurion University of the Negev, Beer-Sheva 8410501, Israel; Russell Berrie Nanotechnology Institute, Technion - Israel Institute of Technology, Haifa 3200003, Israel

## Abstract

The design-build-test (DBT) cycle in synthetic biology is considered to be a major bottleneck for progress in the field. The emergence of high-throughput experimental techniques, such as oligo libraries (OLs), combined with machine learning (ML) algorithms, provide the ingredients for a potential “big-data” solution that can generate a sufficient predictive capability to overcome the DBT bottleneck. In this work, we apply the OL-ML approach to the design of RNA cassettes used in gene editing and RNA tracking systems. RNA cassettes are typically made of repetitive hairpins, therefore hindering their retention, synthesis, and functionality. Here, we carried out a high-throughput OL-based experiment to generate thousands of new binding sites for the phage coat proteins of bacteriophages MS2 (MCP), PP7 (PCP), and Qβ (QCP). We then applied a neural network to vastly expand this space of binding sites to millions of additional predicted sites, which allowed us to identify the structural and sequence features that are critical for the binding of each RBP. To verify our approach, we designed new non-repetitive binding site cassettes and tested their functionality in U2OS mammalian cells. We found that all our cassettes exhibited multiple trackable puncta. Additionally, we designed and verified two additional cassettes, the first containing sites that can bind both PCP and QCP, and the second with sites that can bind either MCP or QCP, allowing for an additional orthogonal channel. Consequently, we provide the scientific community with a novel resource for rapidly creating functional non-repetitive binding site cassettes using one or more of three phage coat proteins with a variety of binding affinities for any application spanning bacteria to mammalian cells.

## Introduction

The design, build, test (DBT) cycle in synthetic biology is considered by many to be a major bottleneck for progress in the field. Specifically, the field is still lacking computational methods that will enable users to reliably design their system of choice without going through multiple time-consuming DBT cycles before arriving at the desired solution. Recent studies are emerging to tackle this challenge and to devise such methods for diverse RNA-, DNA- and protein-based applications, with varying degrees of success^1–3^. Perhaps the best known methods are the Cello algorithm and RBS calculator, which are limited to bacterial chassis^4,5^ at the present time.

The challenge of formulating such algorithms is rooted in the different molecular mechanisms, each governed by diverse dynamics, which in turn require different assumptions and parameter values to model properly using a kinetic or thermodynamic modelling approach. For example, computing the efficiency of ribosome binding sites (i.e. the RBS calculator) requires both a quantitative understanding of ribosome-RNA interactions and RNA structure formation at the translation initiation site. Alternatively, the evaluation of gene regulatory circuits (i.e. the Cello algorithm) requires an understanding of the kinetics of transcription factors binding to DNA, dynamics of mRNA transport, and kinetics of translational processes. Given the particular molecular characteristics needed for each model, it is therefore improbable to generalize these approaches to even related molecular systems (e.g. predicting ribosome binding in mammalian cells). Consequently, since knowledge of the necessary biological parameters is often lacking due to experimental restrictions, kinetic and thermodynamic models are largely qualitative for the vast majority of molecular systems with limited predictive power.

Reliable algorithms are especially needed for the design of RNA-centric functional modules for various applications. A notable study has been recently published, where the authors were successful in designing non-repetitive sgRNA cassettes for targeting multiple metabolic genes in bacteria^6^ demonstrating the utility of such an approach to studying whole systems of genes. Another RNA-based system where a reliable design algorithm can help bring about the full potential of the technology is the encoding of multiple repeats of phage coat protein binding elements on an RNA molecule of choice. In particular, the latter technology has been utilized in many studies and proven to be an important tool in gene editing and RNA-tracking related applications^7–12^. However, current cassettes are characterized by repeated sequence elements, making them nearly impossible to synthesize, clone, and maintain, which limits their utility for robust quantitative measurements^13^ as well as expansion to more complex multi-genic applications.

Previous findings have determined that specificity in phage coat protein binding to RNA is determined by the structural elements formed by specific sequence motifs^14–20^. This implies that for a given phage coat protein, many different sequences may fold into a common functional structure. The DBT problem for phage coat protein-binding cassette design can thus be solved by generating a database of functional binding sites that are divergent from a sequence perspective, and then utilizing different sequences with the same functional structure in place of multiple repeats of the same RNA sequence. The emergence in recent years of high-throughput oligo library (OL) based-experiments^3,21–24^ provides a platform for testing hundreds of thousands of potential binding-site variants. While extremely useful for identifying functional variants, the OL scale is much smaller than the available sequence space for ∼20nt-long binding sites, and thus many functional variants are not sampled. Recently-developed machine-learning (ML) algorithms^25–27^ provide the necessary tool for computationally expanding the variant database to millions of potentially functional sequences, using the OL as an empirical training dataset. The result is an ML algorithm which can computationally score any sequence for the desired functionality

In this work, we apply the OL-ML approach to the design of phage coat protein binding cassettes made of repeated hairpin structures. We generate an OL of many candidate RNA binding sites for the phage-coat proteins MS2, PP7, and Qβ, that are designed to maintain functional RNA structure, despite changes in sequence. We evaluate the function of the resulting RNA hairpins in a massively-parallel *in vivo* expression assay, in bacteria, and subsequently utilize ML tools trained on the OL sequences and their experimental function scores to computational discover and experimentally verify novel sequences that can bind the phage coat proteins with high affinity. Consequently, we demonstrate that sequences with non-repeating elements can be reliably designed, synthesized, and cloned, and, once transcribed, exhibit the functionality expected from the original repeated hairpins, in both bacterial and mammalian cells. This achievement enables researchers to rapidly design functional customized cassettes for RNA-based applications in any organism, effectively eliminating the DBT bottleneck for this technology.

## Results

### Induction-based Sort-Seq (iSort-Seq)

We previously showed that placing a hairpin in the ribosomal initiation region of bacteria can lead to a ∼x10-100 fold repression effect when bound to an RNA-binding protein (RBP)^15,28^. The size of the effect allowed us to adapt this *in vivo* binding assay to a high-throughput OL experiment. We designed 10,000 mutated versions of the single native binding sites to the phage coat proteins of PP7, MS2 and Qβ, and positioned each site at two positions within the ribosomal initiation region (Figure 1A top). The library consists of three sub-libraries within the original library: binding sites that mostly resemble either the MS2-wt site, the PP7-wt site, or the Qβ-wt site (Figure 1A bottom and Figure S1). We introduced semi-random mutations, both structure-altering and structure-preserving, as well as deliberate mutations at positions which previous studies have shown to be crucial for binding. Additionally, we incorporated into our library several dozens of control variants. We used variants characterized in our previous study as positive and negative controls^15,16,29,30^ as follows: positive controls are binding sites that exhibited a strong fold-repression response, and negative control variants are either random sequences or hairpins which did not exhibit a fold-repression response. For the complete library, see Table S1.

**Figure 1:**
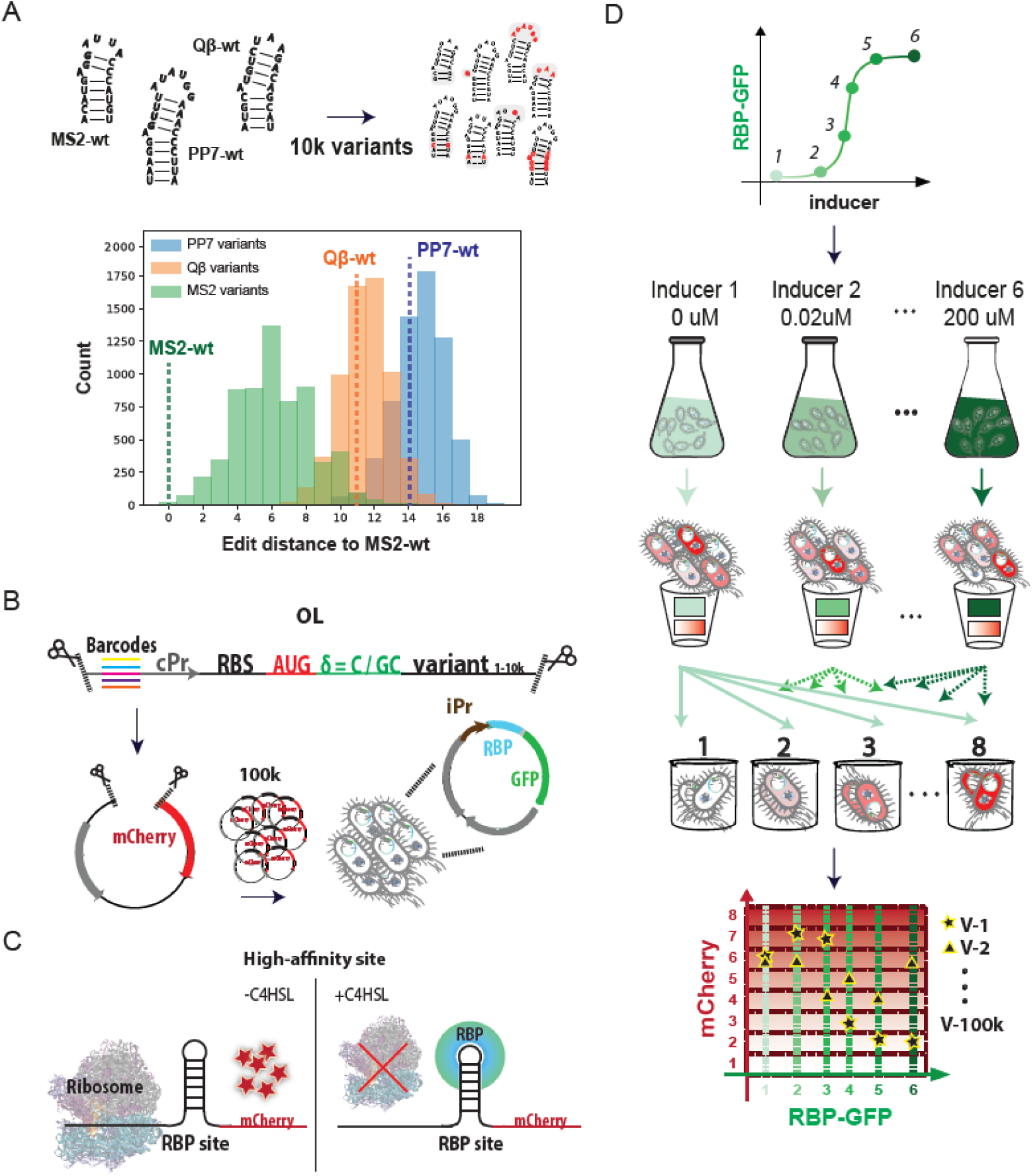
iSort-Seq overview in *E. coli*. (A) (Top) Wild-type binding sites for MS2, PP7 and Qβ phage coat proteins and illustrations of the 20k mutated variants created based on their sequences. (Bottom) Composition of the OL library. Histogram of the number of PP7-based variants (blue), Qβ-based variants (orange), and MS2-based variants (green) with different edit distances from the MS2-wt binding site. (B) Each putative binding site variant was encoded on a 210bp oligo containing the following components: restriction site, barcode, constitutive promoter (cPr), ribosome binding site (RBS), mCherry start codon, one or two bases (denoted by δ), the sequence of the variant tested, and the second restriction site. Each configuration was encoded with five different barcodes, resulting in a total of 100k different OL variants. The OL was then cloned into a vector and transformed into an *E. coli* strain expressing one of three RBP-GFP fusions under an inducible promoter (iPr). The transformation was repeated for all 3 fusions. (C) The schema illustrates the behavior of a high-affinity strain: when no inducer is added, mCherry is expressed at a certain basal level that depends on the mRNA structure and sequence. When inducer (C4HSL) is added, the RBP binds the mRNA and blocks the ribosome from mCherry translation, resulting in a down-regulatory response as a function of inducer concentration. (D) The experimental flow for iSort-Seq. Each library is grown at 6 different inducer concentrations, and sorted into eight bins with varying mCherry levels and constant RBP-GFP levels. This yields a 6×8 matrix of mCherry levels for each variant at each induction level. (Bottom) An illustration of the experimental output of a high-affinity strain (V1) and a no-affinity strain (V2).

We incorporated each of the designed 10k single binding-site variants downstream to an mCherry start codon (Figure 1B) at each of the two positions (spacers δ=C or δ=GC) to ensure high basal expression and enable detection of a down-regulatory response, resulting in 20k different OL variants. Each variant was ordered with five different barcodes, resulting in a total of 100k different OL sequences. The second component of our system included a fusion of one of three phage coat proteins to GFP (Figure 1B) under the control of an inducible promoter. Thus, we created three libraries in *E. coli* cells; each with a different RBP but the same 100k binding site variants. In order to characterize the dose response of our variants, each library was first separated to six exponentially expanding cultures grown in the presence of one of six inducer concentration for RBP-GFP fusion induction. If the RBP was able to bind a particular variant, a strong fold-repression effect ensued, resulting in a reduced fluorescent expression profile (Fig. 1C). We sorted each inducer-concentration culture into eight predefined fluorescence bins, which resulted in an 6×8 fluorescence matrix for each variant, corresponding to its dose-response behavior. We call this adaptation of Sort-Seq “induction Sort-Seq” (iSort-seq - for details see Methods). As an example, we show a high-affinity, down-regulatory dose-response for a positive variant (Figure 1D-bottom V1), and a no-affinity variant exhibiting no apparent regulatory effect as a function of induction (Figure 1D-bottom V2).

### Calculating Binding Scores

We conducted preliminary analysis of the sequencing data to acquire mCherry levels per RBP and inducer concentration, for each variant (Extended Data Figure 2 and SI). We proceeded only with variants that were covered by a significant amount of reads (above 200 reads), and high mCherry basal levels (see SI for further details), for which there is better potential for observing a repression effect. Next, to ascertain the validity of our assay, we characterized the behavior of our control variants (Figure 2A). A linear-like down-regulatory effect as a function of RBP induction is observed for the positive control variants (green), while no response in mCherry levels is observed for the negative controls (red). Additionally, the spread in mCherry at high induction levels is significantly smaller for the positive control than that of the negative control variants. We characterized each variant of the positive and negative controls using a vector composed of three components: the goodness of fit (R-square), the slope of a linear regression, and standard deviation of the fluorescence value at the three highest induction bins (Figure 2B-middle).

**Figure 2.**
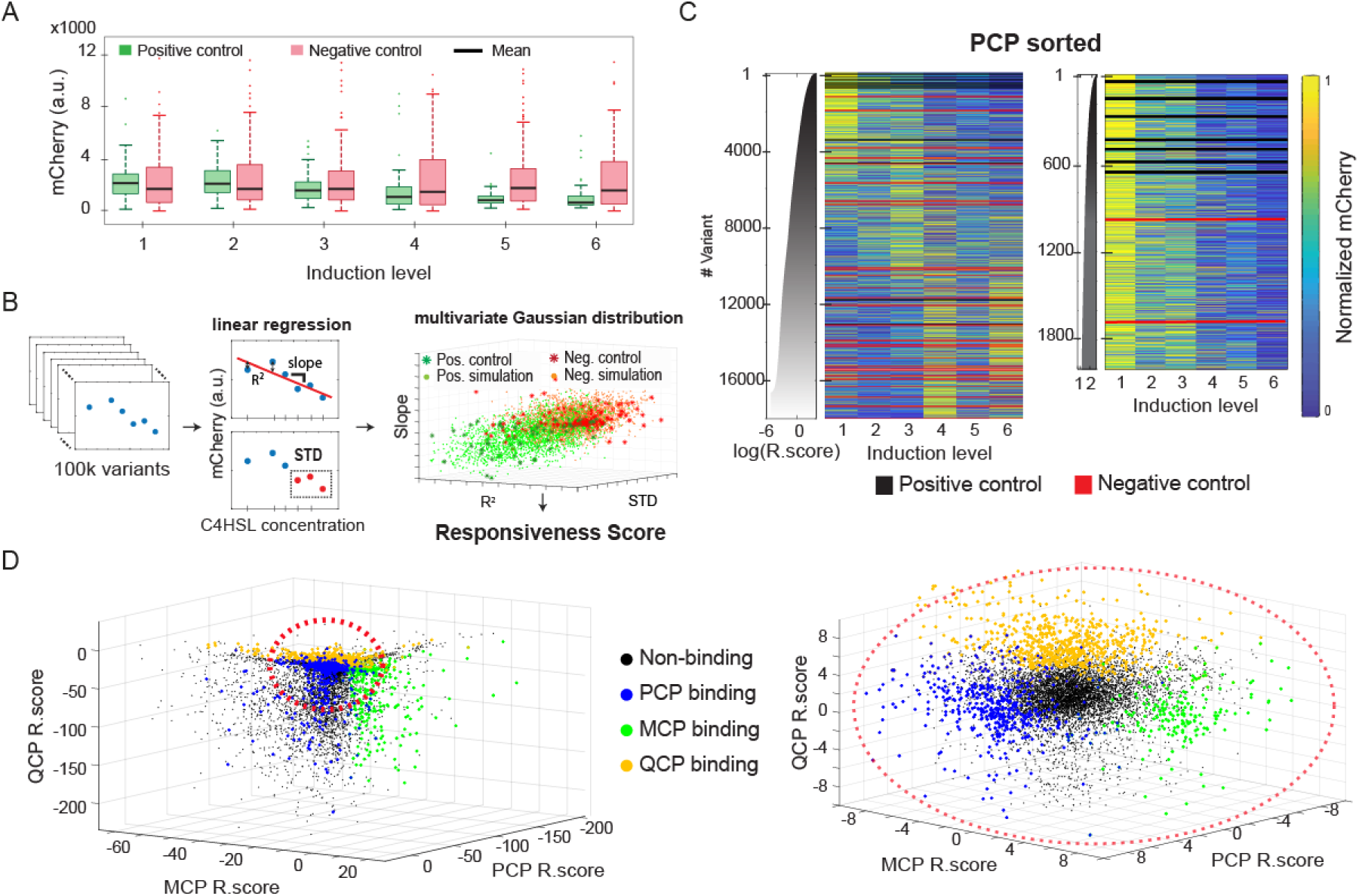
Responsiveness analysis and results. (A) Boxplots of mCherry levels for the positive and negative control variants at each of the six induction levels for PCP-GFP. (B) Schema for Responsiveness score (*R*_*score*_) analysis. (Left & middle) Linear regression was conducted for each of the 100k variants, and two parameters were extracted: slope and goodness of fit (R^2^). The third parameter is the standard deviation (STD) of the fluorescence values at the three highest induction levels. (Right) Location of the positive control (dark green stars) and negative control (red stars) in the 3D-space spanned by the three parameters. Both populations (positive and negative) were fitted to 3D-Gaussians, and simulated data points were sampled from their probability density functions (pdfs) (orange for negative and green for positive). Based on these pdfs the *R*_*score*_ was calculated. (C) (Left) Heatmap of normalized mCherry expression (see SI) for the ∼20k variants with PCP. Variants are sorted by *R*_*score*_. Black and red lines are positive and negative controls, respectively, and the grey graph is the *R*_*score*_ as a function of variant. (Right) “Zoom-in” on the 2,000 top-*R*_*score*_ binding sites for PCP. (D) (Left) 3D-representation of the *R*_*score*_ for every binding site in the library and all RBPs. Responsive binding sites, i.e. sites with *R*_*score*_>3.5, are colored red for PCP, green for MCP, and orange for QCP. (Right) “Zoom-in” on the central highly concentrated region.

Next, we identified subsets of the non-control RNA sequences that were either highly responsive to each RBP (i.e. binding sites with high effective binding affinity), or non-responsive sequences (i.e. sites with no effective binding affinity) based on the positive and negative controls. We computed two multivariate Gaussian distributions using the empirical 3-component vectors that were extracted for the negative and positive controls and for the given RBP, to yield a probability distribution function (pdf) for both the responsive and non-responsive variants (Fig. 2B-right). The two populations are relatively well-separated from one another, presenting two only-slightly overlapping clusters. Finally, we defined the “Responsiveness score” for each variant (*R*_*score*_ – see methods for formal definition) as the logarithm of the ratio of the probabilities computed by the responsive pdf to the non-responsive pdf. This score was computed for each unique barcode, and the final result for sequence variant was averaged over up to five vectors (one for every variant barcode that passes the read-number and basal-level thresholds).

In Figure 2C left, we plot the expression heatmap of the ∼18k variants with PCP (i.e variants that exhibited more than 200 read), sorted (top to bottom) by decreasing *R*_*score*_ (see also Extended Data Figure 3 for MCP and QCP). The plot shows that 5470 variants (counted from the top) exhibit an apparent down-regulatory response, defined as log(*R*_*score*_)>0, corresponding to having a larger probability to belonging to the positive control distribution as compared with the negative. By comparison (Extended Data Figure 3), MCP and QCP yielded 2604 and 7306 such variants, respectively. This indicates that while QCP may be the most promiscuous RBP in our library (i.e. tolerates a more varied set of binding sites), MCP is likely to be the most limited in terms of binding specificity. A closer observation of the top of the list (top 2000, Figure 2C-right) indicates that for a high *R*_*score*_, a rapid reduction in fluorescence is detected in the second bin, which indicates that these variants also seem to exhibit the strongest binding affinity. We next plot the *R*_*score*_ obtained for all three RBPs, for each variant (Fig. 3D). We overlay the plot with colored dots corresponding to the variants with *R*_*score*_ > 3.5 in each list, corresponding to the most specific variants. The plots reveal very little overlap between the subsets of variants that are highly responsive to the different RBPs, indicating that the vast majority of these highly-responsive binding sites are orthogonal (i.e. respond to only one RBP), which was expected for PCP & MCP and PCP & QCP, but not necessarily for MCP & QCP whose native sites are not mutually orthogonal^15–17,20,31,32^.

**Figure 3.**
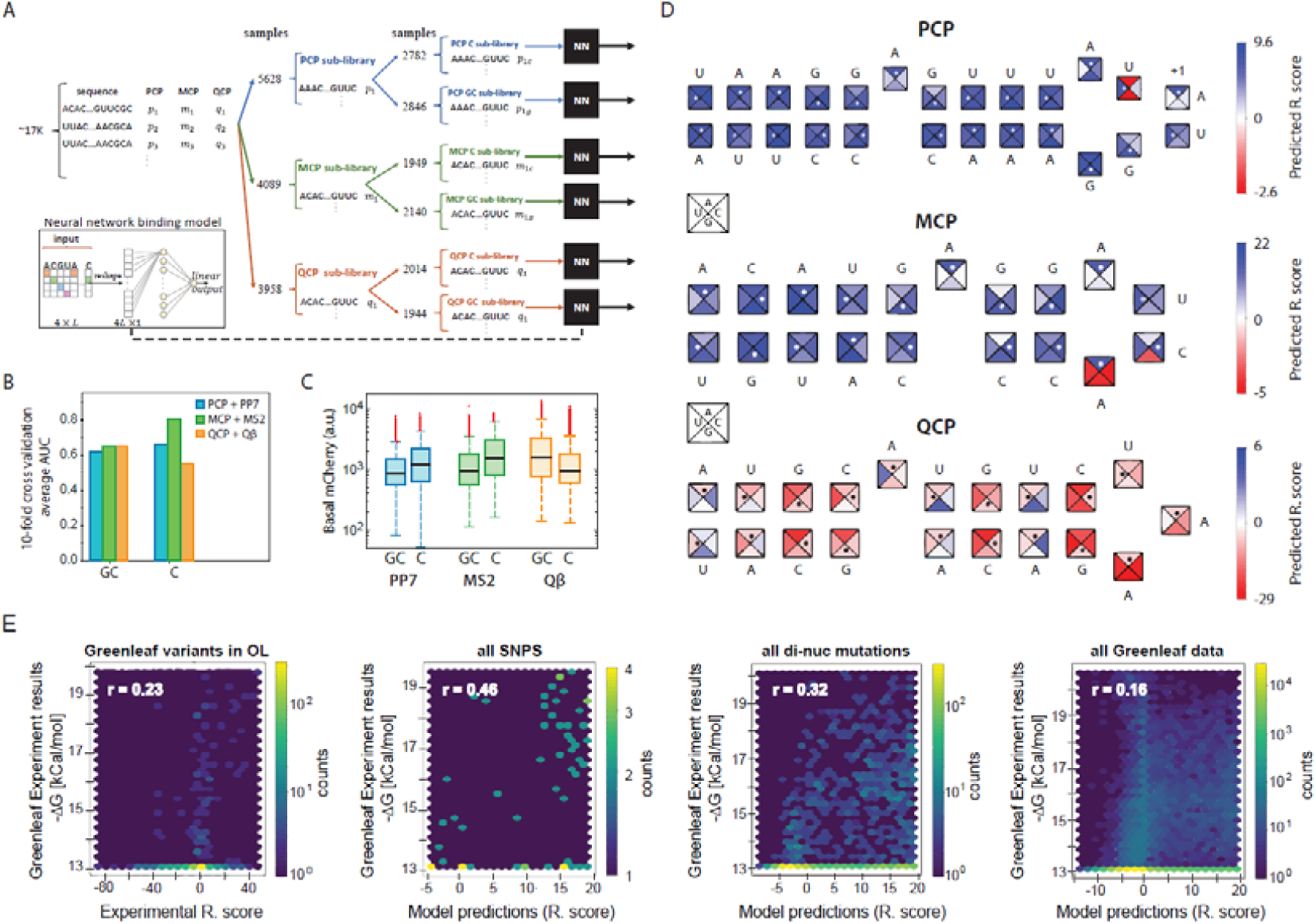
Analysis of MCP, PCP, and QCP RNA-binding preferences. (A) Scheme for the data preparation and neural network (NN) architecture (inset) used. (B-C) Average AUC of the 10-fold cross validation (B) and box plots of the mCherry basal levels (C) conducted on the six sub-libraries: PCP with PP7-based binding sites, MCP with MS2-based binding sites, and QCP with Qβ-based binding sites, all with either δ=GC or δ=C. (D) Illustrations of the NN predictions for the three sub-libraries for any single-point mutation. Each binding site is shown, with the wild-type sequence indicated as letters above and white dots inside the squares. Each square is divided to the four possible options of nucleotide identity, with the colors representing the predicted *R*_*score*_ for each option. (E) *R*_*score*_ comparison to δG results of a previous study that reported MCP binding to more than 129k sequences^19^. Each plot (from left-to-right) represents the correlation coefficient using: the experimental measurements for variants that were both in our OL and in the *in vitro* study, the *R*_*score*_ values predicted by our ML model for all single-mutation variants, for all double-mutation variants, and for the entire set of 129,248 mutated variants.

### Computational analysis of RNA-binding preferences

Using empirical *R*_*score*_ values and associated binding site sequences as training set, we developed an ML-based method that predicts the *R*_*score*_ values for every mutation in the wild type (WT) sequences. We first built a model specific to each protein and its WT length to validate our OL measurements on prior knowledge of the proteins’ binding specificities. To do so, we used a neural network, that receives as input the sequence of a binding site and outputs a single score. We train a specific network for each of the three RBP-OL experiments (Figure 3A and SI). The network only considers variants with the same length as the three WT sequences (25nt for PP7-wt, 19nt for MS2-wt, and 20nt for Qβ-wt), and all variants were split into two categories according to their position within the ribosomal initiation region (Figures 3A and Extended Data Figure 4), resulting in a total of six different models. Such a model preserves the positional information for each feature, i.e. the position of each nucleotide in the WT binding site. To choose the position (spacer δ) in which more robust scores were measured, we looked at the average AUC (area under the receiver operating characteristic curve) over 10-fold cross validation. The AUC scores for the most robust position yielded values of 0.65 for PCP with PCP-based sites with δ=C, 0.8 for MCP with MCP-based sites with δ=C, and 0.65 for QCP with QCP-based sites with δ=GC (Figure 3B). Interestingly, the variant group with higher AUC score was also characterized by higher basal mCherry expression levels (Figure 3C), which in turn resulted in a higher fold repression effect. Thus, higher AUC, meaning more robust predictability, correlated with higher fold-repression, which provided additional validity to our analysis.

**Figure 4.**
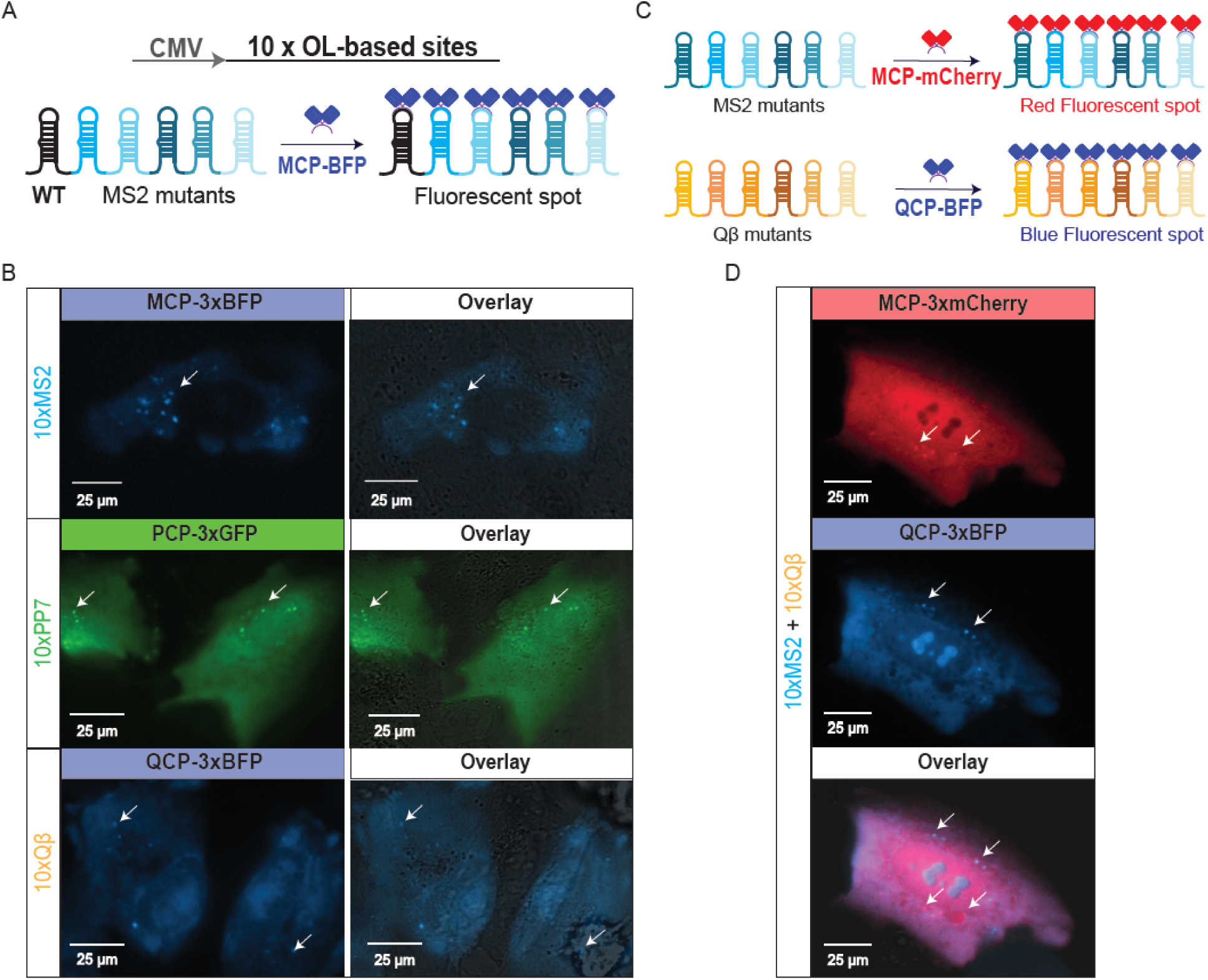
Universal cassettes for RNA imaging in U2OS cells. (A) Experiment design for the three cassettes based on the experimental binding sites. High *R*_*score*_ binding sites were incorporated into a ten-site cassette downstream to a CMV promoter. When the matching RBP-3xFP is added (MCP-3xBFP is shown), it binds the binding-site cassette and creates a fluorescent spot. (B) The results for all three cassettes transfected with the matching RBP-3xFP plasmid into U2OS cells and imaged by fluorescence microscopy for detection of fluorescent *foci*. For each experiment, both the relevant fluorescent channel and the merged images with the differential interference contrast (DIC) channel are presented. (C) Experimental design for the orthogonality experiment: two separate cassettes with 10 predicted dual-binding, mutated (no-WT) sites for only MCP or only QCP, respectively were designed and transfected together with both MCP-3xmCherry and QCP-3xBFP, into U2OS cells. (D) Results for the orthogonality experiment: a cell presenting non-overlapping fluorescent *foci* from both fluorescent channels, indicating binding of MCP and QCP to different targets. Fluorescent wavelengths used in these experiments are: 400nm for BFP, 490nm for GFP, and 585nm for mCherry.

In order to better understand the relationship between binding site sequence and binding, we used the model to analyze the effect of single-point mutations in each of the WT binding-site sequences, for each of the OL-RBPs (Figure 3D). We present the ML model’s results as “binding rules” depicted in illustrations for each of the three RBPs. The schemas represent the predicted responsiveness for every single-nucleotide mutation to the WT sequences. For instance, in the schema for PCP, mutating the bulge from A to C or U sharply reduces the structure’s predicted responsiveness. In addition, mutation of the second nucleotide in the loop from U to either A or G quickly abolishes the predicted responsiveness, while mutation of the sixth nucleotide in the loop leaves binding unaffected. A clear characteristic of PCP is the tolerance to single nucleotide mutations, which is reflected by the dominance of the blue colors for most mutations in the lower stem. The PCP binding site also appears to be tolerant to mutations in the upper stem, while the loop region (particularly the first three nucleotides) appear to be highly sensitive to mutations. It is important to note that our results for PCP broadly correlate with past works^15,16,18^, which found that loop region and bulge to be critical for PCP binding.

For MCP, a tolerance to mutations in the lower stem emerges from our analysis, while a strong sensitivity to mutations in the bulge and the loop regions is revealed. Past analysis^15,19,30^ also highlighted the sensitivity to mutations in the loop and bulge regions, indicating that the *in vivo* environment does not alter the overall binding characteristics of MCP. Finally, for QCP (Fig. 3D-bottom), a significantly different picture emerges, as it seems that the native sequence we used (as referred to in the literature^14,15,17^) is far from the highest-affinity sequence for binding. In most cases, it has a lower *R*_*score*_ than the mutated versions. The bulge, for instance, has a much higher *R*_*score*_ with U instead of the native A. Consequently, our data seems to indicate that QCP prefers a GC-rich stem and a U bulge, which is structurally different than the bulge-stem-loop and stem-loop preferences of MCP and PCP, respectively. Altogether, the three proteins preferably bind different motifs and structural elements, which explains the orthogonality observed in our data (Figure 2D).

To validate both our experimental measurements and model predictions, we compared our results to a previous study that measured high-throughput *in vitro* RNA-binding of MCP^19^. In the study, the researchers employed a combined high-throughput sequencing and single molecule approach to quantitatively measure binding affinities and dissociation constants of MCP to more than 10^7^ RNA sites using a flow-cell and *in vitro* transcription. The study reported Δ*G* values for over 120k variants, which formed a rich dataset to test correlation with our measured and predicted *R*_*score*_ values. First, we computed Pearson correlation coefficient using the purely experimental measurements for variants that were both in our library and in the *in vitro* study. The result (Figure 3E-left) indicates a positive and statistically significant correlation (R=0.23). We next predicted *R*_*score*_ values using our model for all the reported variants of the *in vitro* study (Figure 3E left-to-right), and found a strong correlation (R=0.46) for single-mutations variants, a moderate correlation (R=0.32) for double-mutation variants and a weak correlation (R=0.16) with the entire set of 129,248 mutated variants. Given the large difference between the experiments and the different sets of variants used (e.g. *in vitro* vs. *in vivo*, microscopy-based vs. flow cytometry-based), the positive correlation coefficients (p-values<0.0002) indicate a good agreement for both sets of experimental data, and a wide applicability for the learned binding models for MCP.

### Universal new cassettes for RNA imaging

To further validate the results of our experiment and test the wider applicability of the findings, we generated new cassettes containing multiple non-repetitive RBP binding sites identified by our experimental data set, and tested them in mammalian cells. Once labelled with a fusion of the RBP to a fluorescent protein, functional cassettes appear as trackable bright fluorescent *foci*. We designed three binding site cassettes based on library variants that were identified as highly responsive for each RBP (Figure 4A). Each cassette was designed with ten different binding sites, all characterized by a large edit distance (i.e. at least 5) from the respective WT site, thus creating a sufficiently non-repeating cassette that IDT was able to synthesize in three working days. In addition, all selected binding sites exhibited non-responsive behavior to the two other RBPs in our experiment. We cloned the cassettes into a vector downstream to a CMV promoter for mammalian expression and transfected them into U2OS cells together with one of the RBP-3xFP plasmid encoding euther PCP-3xGFP, MCP-3xBFP, or QCP-3xBFP. In a typical cell (Figure 4B), all three cassettes generated more than five fluorescent puncta, dispersed throughout the cytoplasm. The puncta were characterized by rapid mobility within the cytoplasm, and a lack of overlap with static granules or distinct features which also appear in the DIC channel (see Supplementary movies).

To expand our claim to orthogonal and simultaneous imaging of multiple promoters, we ordered two additional cassettes with MS2 and Qβ variants, respectively, and co-transfected them with a plasmid encoding for both of the matching fusion proteins: MCP-3xmCherry and QCP-3xBFP (Figure 4C). For each cassette, the sites were chosen with two constraints: to minimize repeat sequences and to maximize orthogonality to the other RBP (e.g. both MS2-wt and Qβ-wt binding sites were not included as they exhibit cross-responsiveness and are thus not orthogonal). In Fig. 4D we plot sample cell images depicting single and double channel views. The images show that both cassettes produce a spatially distinct set of puncta (Fig. 4D-top and middle), which can be definitively associated with one of the two proteins (Fig. 4D-bottom). This indicates that our binding sites are sufficiently orthogonal to allow tracking of more than one cassette simultaneously. Moreover, there is little difference between the number of puncta of the two sequences and the fluorescent intensity for all puncta seem to fluctuate unimpeded in all three directions (x,y, and z) inside the cell. Taken together, the microscopy experiments conducted in mammalian cells demonstrate the universal applicability of the results obtained from the high-responsiveness binding sites identified in the OL experiment to the advancement of RNA imaging in a variety of cell types.

### *De novo* design of dual-binding site cassettes

Finally, to further verify the validity of the ML-approach, we opted to test the predictive power of the model. To do so, we altered our initial ML approach by developing a protein-specific model based on the whole library, which we termed whole-library model (Figure 5A, Extended Data Figure 5 for AUC curves, and further details in the SI). This model, as opposed to our NN, enables binding prediction to any site, i.e. of length different than the WT-site length. The model is based on a convolutional neural network (CNN). It receives as input both the sequence and structure of the RNA binding site, as calculated by RNAfold^33^. It is composed of one convolution layer, a pooling layer, a fully connected layer and a single output node. All three CNNs showed improved predictive capabilities when the structural data was added into the network (Extended Data Figure 6).

**Figure 5.**
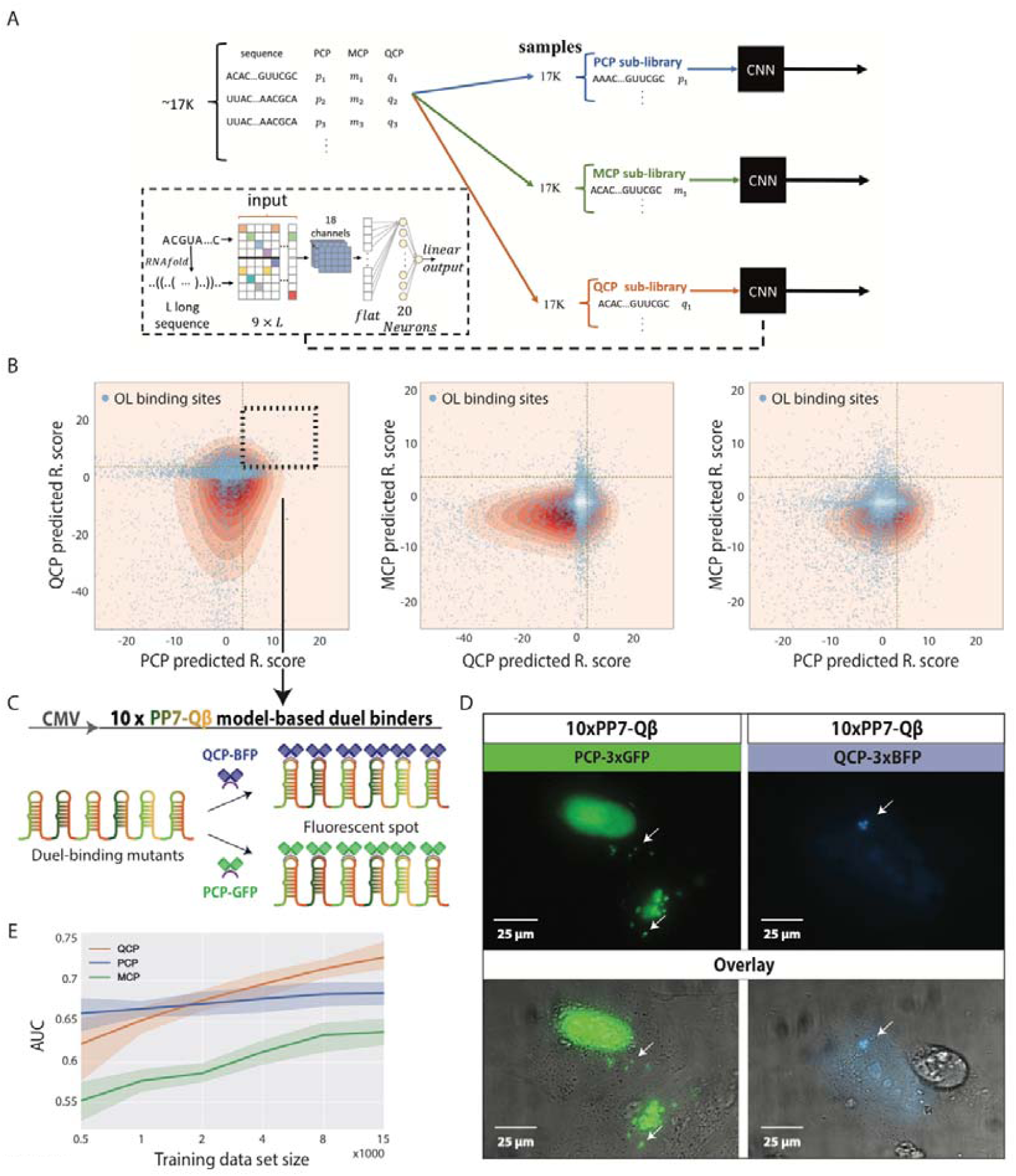
*De novo* design of dual-binding site cassettes in U2OS cells. (A) A scheme for the data preparation and neural network (NN) architecture (inset) used for the protein-specific convolutional neural network (CNN) model based on the whole library. (B) 2D density plots (pink-red scale) depicting the predicted *R*_*score*_ values for one million ML variants binding to (left-to-right): PCP and QCP, MCP and QCP, and MCP and PCP. QCP-PCP dual-binding variants are located in the black dashed square. Blue-white dots represent the experimental OL variants. (C) Based on the dual-binding mutants for QCP and PCP, we designed an additional cassette based on our models’ predictions alone (no binding sites from the original OL). (D) Results for the dual-binding experiment. Fluorescent *foci* can be observed for the cassette expressed with either PCP-3xGFP or QCP-3xBFP. For both experiments, both the relevant fluorescent channel and the merged images with the DIC channel are presented. Fluorescent wavelengths used in these experiments are: 400nm for BFP and 490nm for GFP. (E) Evaluation of prediction accuracy based on size of the training set. For each training set size, a random set of more than 1,000 training-set variants was withheld for computational testing post-training. Performance is reported as average AUC over 10 random training and test sets (and standard deviation in shade).

We used the whole-library model to predict *de novo* functional binding site sequences, which could bind multiple RBPs. To do so, we generated all possible variants with Hamming distance 3-7 to one of the three WTs. From this set of sequences, we randomly selected one million sequences and used the models to predict the responsiveness score for each of the three RBPs. In Figure 5B, we plot the variant density distribution based on a predicted *R*_*score*_ values. The plots show that the highest density of sequences appears at *R*_*score*_ values that hover around 0 for all three proteins. The plots further show that there is a bias towards negative responsiveness values for all three proteins in the computed sequences. This is consistent with having a small region of sequence space which facilitates specific binding, which in turn is easy to abolish with a small number of mutations. In contrast, high responsiveness scores are only computed for a small number of the sequences, as can be seen by the sharp gradient in the density plot for positive responsiveness values. Finally, each plot shows a non-negligible region where the same sequence exhibits a high responsiveness score for both RBPs. These sequences are predicted to be double binders. By overlaying the empirical responsiveness score for all the variants in our library (white and blue dots), we observe that the dual-binder region is inhabited by a handful of experimental variants for each possible RBP pair.

To test the predictions of the whole-library models experimentally, we designed *de novo* cassette sequences that consisted only of predicted dual-binder variants, with orthogonal WT sites. We designed another 10x binding site cassette (Fig. 5C), where each binding site was selected from the set of predicted sequences whose responsiveness scores for QCP and PCP were both above 3.5 (see dashed square in Fig. 5B-left panel). Therefore, we expected the cassette to generate fluorescent *foci* when bound by either QCP or PCP. As before, we cloned the cassette into a vector downstream of a CMV promoter for mammalian expression and transfected it into U2OS cells together with a plasmid encoding for either PCP-3xGFP or QCP-3xBFP. In Figure 5D, we plot fluorescent and DIC images for PCP (left) and QCP (right), depicting bright fluorescent *foci* that are located outside of the nucleus and which do not overlap with a DIC feature. The plots show distinct puncta observed with both relevant RBPs confirming the dual binding nature of the cassette. Consequently, these images support our model’s ability to accurately predict MCP, PCP, and QCP binding sequences with known function with respect to all three RBPs.

## Discussion

In this study, we adapted our previously-developed binding assay for quantifying RBP-RNA binding site affinity *in vivo* to a high-throughput OL platform in bacteria^15,28^. Consequently, we were able to generate a sufficiently large dataset for training an ML-based model to reliably predict functional non-repeating binding elements for the phage coat proteins of PP7, MS2, and Qβ. As a result, we now possess a computational tool that allows us to bypass the DBT-cycle when designing new molecules encoding multiple repeats of these binding sites. This tool substantially shortens the time from design to functional applications and removes many of the previous restrictions associated with these systems, such as the need for repetitive cloning cycles, repeat-based structure formation, and limitation on the number of functional binding sites. We also demonstrated that our MCP and QCP sites are orthogonal to one another, allowing for an additional orthogonal channel. These achievements provide the community with a reliable design tool for new phage coat protein binding cassettes in a variety of organisms.

In addition to solving an important technological bottleneck, we were also inspired by the need to develop new approaches for understanding RNA-related problems. It is generally believed that the combinatorial nature of RNA sequence and its intramolecular interactions lead to high complexity, making simulations based on biophysical models a difficult task with limited degree of success^33–36^, even when cellular environment is not taken into account. As a result, little is known about the evolutionary constraints on RNA structures, making bioinformatic identification of functional RNAs difficult^19^. In this work, using the OL-ML approach, we were able to quantitatively model *in vivo* binding of three phage coat proteins to RNA, at single-base-pair resolution, and for every possible single-nucleotide mutation. Based on the model, we concluded that the binding site for QCP is not optimal, and we could design *de novo* dual-binding binding sites (PCP *and* QCP) as well as orthogonal binding sites (MCP*-only* or *Q*CP-*only*) that did not exist naturally. For such an endeavor to work, we had to achieve a level of understanding beyond that accomplished by typical single-clone approaches. Furthermore, our demonstration that modeling of single RNA binding sites in bacteria is sufficient for generating a reliably predictive model for multi-binding site cassettes in mammalian cells is evidence that, at least for this set of proteins, the RNA-RBP module can accommodate multiple cellular environments, thus constraining the complexity of the overall system. Consequently, our work paves the way for characterizing and predicting binding of additional RBPs in any cellular environment, in addition to providing a proof-of-concept for the OL-ML approach.

Finally, our work not only provides a blueprint for studying RNA-related systems, but also partially answers the question of how much data is needed to train a reliably predictive model that will allow one to bypass the DBT-cycle. In our case, several thousand variants were sufficient (Fig. 5E). At the present time, it is impossible to tell whether this number is typical or “surprisingly” small, as there are very few experiments to compare with. However, given our previous (albeit partial) mechanistic and structural understanding regarding PCP, MCP, and QCP binding to RNA that informed the OL design process, a reduction in the number of OL variants needed for the learning process was expected. It is reasonable to assume that partial knowledge of a system could reduce the size of the useful training set, and is likely to be an important ingredient in generating a complete computational understanding of a system. Future work on other complex RNA-based molecular interaction systems will determine whether the OL-ML approach is indeed a useful tool for providing new mechanistic and structural insights into these systems.

## Methods

### Bacterial OL work

#### Construction of the bacterial library

We designed 10,000 mutated versions of the WT binding sites to the phage coat proteins of PP7 (Figure 1 and Extended Data Fig. 1), MS2 and Qβ, and positioned them at two positions within the ribosomal initiation region. Each of the designed 10k sites were positioned either one or two nucleotides downstream to the mCherry start codon, resulting in 20k different configurations. We then ordered the following oligo library (OL) from Agilent: 100k oligos (Extended Data Table 1), each 210bp long containing the following components: BamHI restriction site, barcode (five for each variant), constitutive promoter (cPr), Ribosome Binding Site (RBS), mCherry start codon, one or two bases (denoted by delta), the tested binding site, ∼60 bases of the mCherry gene, and an ApaLI restriction site. We then cloned the OL using restriction-based cloning strategy. Briefly, the 100k-variant ssDNA library from Agilent was amplified in a 96-well plate using PCR (see Extended Data Table 2 for primers), purified, and merged into one tube. Following purification, dsDNA was cut using BamHI-hf and ApaLI and cleaned. Resulting DNA fragments were ligated to the target plasmid containing an mCherry open reading frame and a terminator, using a 1:1 ratio. Ligated plasmids were transformed to E. cloni® cells (Lucigen) and plated on 37 large agar plates with Kanamycin antibiotics in order to conserve library complexity. Approximately two million colonies were scraped and transferred to an Erlenmeyer for growth. After O/N growth, plasmids were extracted using a maxiprep kit (Agilent), their concentration was measured, and they were stored in an Eppendorf tube in −20.

#### Construction of RBP-GFP fusions

RBP sequences lacking a stop codon were amplified via PCR of either Addgene or custom-ordered templates (Genescript or IDT, see Extended Data Table 3). MCP, PCP and QCP were cloned into the RBP plasmid between restriction sites KpnI and AgeI, immediately upstream of a GFP gene lacking a start codon, under the pRhlR promoter (containing the rhlAB las box38) and induced by C4-HSL. The backbone contained an Ampicillin (Amp) resistance gene. The resulting fusion-RBP plasmids were transformed into *E.coli* TOP10 cells. After Sanger sequencing, positive transformants were made chemically competent and stored at −80°C in 96-well format.

#### Double Transformation of OL and RBP-GFP plasmids

Note: steps 3 to 5 were conducted three times, one for each RBP-GFP fusions.

OL DNA was transformed into ∼300 chemically competent bacterial cell in 100ul aliquots containing one of the RBP-mCeulean plasmids in 96-well format. After transformation, cells were grown in 2L liquid LB with twice the concentration of the antibiotics – Kanamycin and Ampicillin – overnight at 37°C and 250rpm. After growth glycerol stocks were made by centrifugation, re-suspension in 30ml LB, mix 1.2ml with 400ul 80% glycerol – 20% LB solution and store in −80°C.

#### Induction-based Sort-Seq OL assay

One full glycerol stock of the library was dissolved in 500ml of LB with antibiotics and grown overnight at 37°C and 250rpm. In the morning, the bacterial solution were diluted 1:50 into 100ml of semi-poor medium consisting of 95% bioassay buffer (BA: for 1L - 0.5g Tryptone [Bacto], 0.3ml Glycerol, 5.8g NaCl, 50ml 1M MgSO4, 1ml 10xPBS buffer pH 7.4, 950ml DDW) and 5% LB. The inducer, C4-HSL, was pipetted manually to a final concentration of one out of six final concentrations: 0uM, 0.02uM, 0.2uM, 2uM, 20uM, and 200uM. Cells were grown at 37°C and 250rpm to mid-log phase (OD600 of ∼0.6) as measured by a spectrophotometer and taken to the FACS for sorting.

During sorting by the FACSAria cell sorter each inducer level culture was sorted into eight bins of increasing mCherry levels spanning the entire fluorescence range except for 5% at the higher end (bin 1 - low mCherry to bin 8 - high mCherry), and set GFP levels (for example, the 0mM culture were sorted according to zero GFP fluorescence, the 0.02uM culture to slightly positive GFP fluorescence, and so on). Sorting was done at a flow rate of ∼20,000 cells per sec. 300k cells were collected in each bin for the entire 6×8 bin matrix. After sorting, the binned bacteria were transferred to 10ml LB+KAN+AMP and shaken at 37°C and 250rpm overnight. In the morning, cells were prepared for sequencing (see below) and glycerol stocks were made by mixing 1ml of bacterial solution with 500ul 80% glycerol – 20% LB solution and stored in −80°C.

#### Sequencing

Cells were lysed (TritonX100 0.1% in 1XTE: 15μl, culture: 5μl, 99°C for 5 min and 30°C for 5 min) and the DNA from each bin was subjected to PCR with a different 5’ primer containing a specific bin-inducer level barcode. PCR products were verified in an electrophoresis gel and cleaned using PCR Clean-Up kit. Equal amounts of DNA (2ng) from 16 bins were joined to one 1.5ml microcentrifuge tube for further analysis, to a total of three tubes. This procedure was conducted three times, one for each RBP-GFP fusions.

Each one of the three samples were sequenced on an Illumina HiSeq 2500 Rapid Reagents V2 50bp 465 single-end chip. 20% PhiX was added as a control. This resulted in ∼540 million reads, about 180 million reads per RBP.

### Mammalian Cassette microscopy experiments

#### Construction of mammalian expression plasmids

We ordered three plasmids from addgene containing PCP-3xGFP (#75385), MCP-3xBFP (#75384), and N22-3xmCherry (#75387), and used them to create the following two plasmids: MCP-3xmCherry and QCP-3xBFP. In brief, using two restriction enzymes, BamHI and Mlul, we restricted the plasmids and conducted PCR with the same restriction sites added as primers on both MCP and QCP. After PCR purification, we restricted the product with the same two enzymes and ligated them to the matching plasmids. Then, we performed transformation to Top10 *E.coli* cells and screened for positive clones. All plasmids used in the microscopy experiments were sequence-verified via Sanger sequencing.

RNA binding site cassettes were ordered from IDT as g-blocks (see Extended Data Table 4 for sequences). Restricted and ligated them to a vector downstream of a CMV promoter using the restriction enzyme EcoRI. Then, we performed transformation to Top10 *E.coli* cells and screened for positive clones. All plasmids used in the microscopy experiments were sequence-verified via Sanger sequencing.

#### Mammalian Microscopy assay

##### 1. Cell culture

The Human Bone Osteosarcoma Epithelial Cell line (U2OS, see Key Resource table) was incubated and maintained in 100×20mm cell culture dishes under standard cell culture conditions at 37°C in humidified atmosphere containing 5% CO_2_ and were passaged at 80-85% confluence. Cells were washed once with 1x PBS, and subsequently treated with 1mL trypsin/EDTA (ethylenediaminetetraacetic acid, Biological Industries) followed by incubation at 37°C for 3-5 minutes. DMEMcomplete, complemented with 10% FBS and final concentrations of 100U penicillin plus 100μg streptomycin, was added and transferred into fresh DMEMcomplete in subcultivation ratios of 1:10.

##### 2. Fluorescent microscopy experiments

Before the experiment, U2OS cells were seeded on 60mm glass-bottom imaging dishes. Transient transfection was performed with Polyjet (XX) transfection reagent according to the manufacture’s instructions. Typical DNA for transfection was 150ng from RBP-3xFP and 850ng from the cassette plasmid. After inculabtion of 24-48 hours, the growth medium was removed and replaced with Leibovitz L15 medium with 10% FBS. During microscopy, the sample was kept at 37°C.

Images were taken with a 40X oil immersion objective and the following excitation lasers: 585nm for mCherry, 490nm for GFP, 400nm for BFP. The images were recorded with an EMCCD camera. The microscope was controlled with NIS Elements imaging software. Time-lapse movies of a single Z-plane were recorded with, 1500ms exposure time and time intervals between frames were 30 seconds.

### Analysis

#### Responsiveness score

Note: the following analysis procedure was conducted three times, one for each RBP.

##### 1. Read normalization and filtration

Read number was normalized by percentage of bacteria in each bin from the total library, given by the FACS during sorting. This is done in order to be able to compare between numbers of reads of the same variant in different bins.

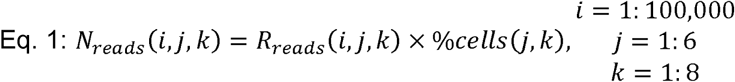

where *N*_*reads*_*(i,j,k)* and *R*_*reads*_ are the number of normalized and raw reads per variant, bin, and inducer concentration respectively. *%cells (j,k)* corresponds to the percentages of the cells of variant *i* in each bin per inducer concentration during sorting from the entire library as supplied by the sorter.

Two cut-offs were introduced on the variant read counts: (i) only inducer levels that had above 30 reads for all eight bins were taken into account; and (ii) only variants that had more than 300 reads in total for the entire 6-by-8 matrix were taken into account.

##### 2. Estimation of mean mCherry levels (μ) per inducer concentration from reads per variant

For each inducer concentration *j*, we have an 8-bin histogram for which we need to calculate the mCherry averaged fluorescence (μ*(i,j)*). First, for every variant we renormalize *N*_*reads*_ by the total number of reads obtained for that inducer level (each column in the read matrix and color bar, Extended Data Figure 2A-top).

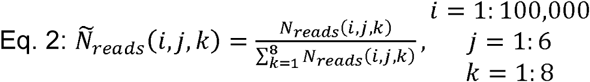

Next, we convert the bin index (j=1:8) to mCherry fluorescence (*Bin(i,j,k)*). This is done by retrieving the maximum mCherry fluorescence value that was assigned to each bin by the sorter. Then, we compute the cumulative renormalized reads by adding all the normalized reads successively from the lowest to the highest fluorescent bin as follows:

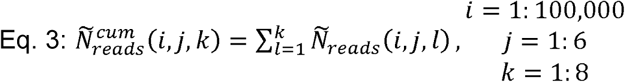

Finally, to compute μ, we fit the cumulative renormalized read values to a cumulative Gaussian as follows:

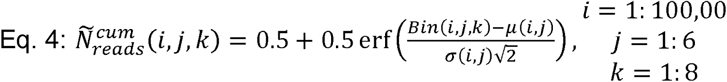

where σ*(i,j) i*s the standard deviation for mCherry fluorescence extracted from the fitting procedure (see Extended Data Figure 2A-bottom for sample calculation). Note, only induction levels that had a goodness-of-fit higher than 0.5 were taken into account in the final analysis.

##### 3. µ normalization and filtration

Since each inducer concentration experiment was carried out in different conditions (e.g. duration of incubation on ice, O/N shaking, binning time) and at a different time (different days), mCherry levels assigned for each bin varied greatly as a function of experiment as well as over-all fluorescence recorded. Therefore, to quantify this systematic error, we first computed a normalized mean fluorescence level (μ_*norm*_) per variant as follows:

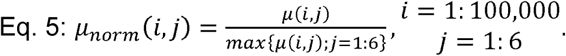

To ascertain the scope of the problem presented by the systematic error, we plot in Extended Data Figure 2B a heat-map of μ_*norm*_ values consisting of 3000 variants for PCP. Here, low fluorescence was recorded for induction level 1, 4, and 6, while higher levels were recorded for induction levels 2,3, and 5 respectively. These results are consistent with the fact that the induction experiments of level 1,4, and 6 were carried out on the same day, while those of 2,3, and 5 on a separate day.

Next, to accommodate for these systematic discrepancies in our data, for each inducer level we extracted the μ_*norm*_ for all the negative control variants that were introduced into the OL (220 variants for PCP, 160 variants for MCP and QCP). We then computed the average μ_*norm*_ for all negative controls per inducer level to obtain μ_*neg*_*(j).* Finally, we rescaled all μ_*norm*_*(i,j)* values by μ_*neg*_*(j)* to eliminate the systematic error from the average fluorescence level as follows:

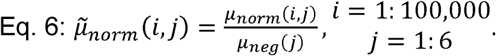

Extended Data Figure 2C shows that this rescaling operation successfully compensated for the systematic error. Note, that since the experiment is based on detecting a repression effect as a function of inducer, we filtered out the variants that displayed averaged mCherry levels at the three lowest concentrations below 15% of the averaged mCherry levels at the three lowest concentrations of the positive control.

##### 4. Calculating the responsiveness score (R_score_)

Next, we mapped each variant’s 6-value average fluorescence vector 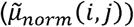 to an empirical or ad-hoc 3-vector of principle components consisting of a *slope (m)* and *goodness-of-fit* (*R*^*2*^) to a simple linear fit of the rescaled fluorescence 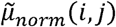 to inducer concentration values. The third principle component is a standard deviation (*std)* of 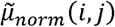 computed at the three highest concentration induction bins. We term this new vector:

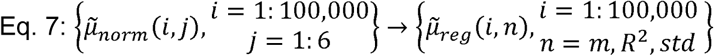

Next, we computed a probability density function for both the positive and negative controls as follows:

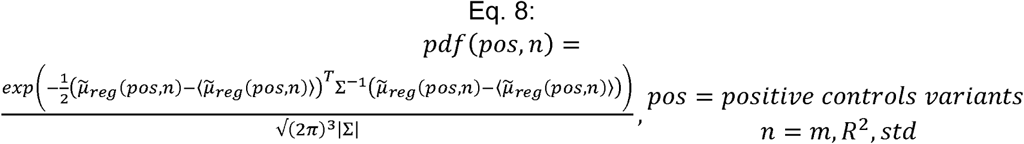

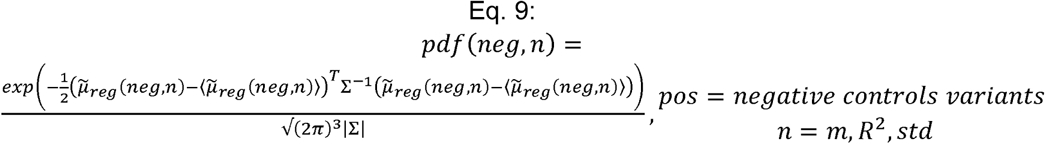

where *pdf(pos,n)* and *pdf(neg,n)* are the probability density functions computed over all the positive and negative control variants respectively, and ∑ is the co-variance matrix (3-by-3).

Using these probability density functions, we then computed the probability that a random variant (*i*) belongs to each of these distributions, as follows:

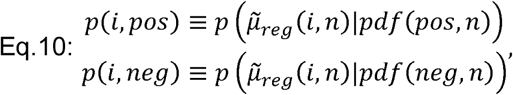

Which then allows us to calculate a responsiveness score (*R*_*score*_), which is a measure that quantifies the likelihood that a particular variant’s 3-vector behaves similarly to those of the positive controls. This observable is defined as follows:

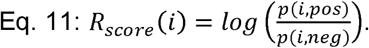

Finally, since every variant in our library appears five times, we averaged the *R*_*score*_ over as many bar-codes which past our filters, and proceed to sort the 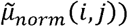 6-vectors in accordance with decreasing *R*_*score*_ in the heatmaps presented in Fig.2 and Extended Data Figure 3.

#### Machine-learning methods

We developed two types of models to predict the binding preferences of the three RNA binding proteins (RBPs): WT-specific and whole-library. Here, we will describe in detail the models, the choice of hyper-parameters and their training on experimental data. First, we will cover the features common to the two models. Then, we will provide details relevant to each of the two model types separately.

##### Dataset

The dataset contains RNA-binding intensity of three proteins (MCP, PCP and QCP) to approximately 17,000 sequences (PCP 17,177, MCP 17,213, QCP 16,041, and 12,245 in the intersection of the three). All sequences were either a variant of a known wild type (WT) binding site of one of the three proteins or a non-similar sequence that was used as control (PCP 42, MCP 40, QCP 38). The edit distance of the derived sequences from their WT mostly span 4 to 8 mutations or indels (Extended Data Figure 1). The binding intensity score, also denoted as the responsiveness score, empirically spanned the range of −281 to 47. Each sequence has a positional feature, which defines its prefix and suffix, i.e. upstream and downstream flanking sequences, respectively. The prefix is either C or GC and the corresponding suffix is one out of three options: T, CT or no suffix. The choice of suffix is done in a way that guarantees no shift in the reading frame.

##### Data encoding

To provide the sequence data as input to the computational framework we used, it first needs to be transformed to numerical values. Each sequence was encoded using a traditional one-hot encoding of the sequence. Each nucleotide is converted to a four-bit vector with one bit set in the position corresponding to that nucleotide and all other positions set to zero. This way an L-long sequence is transformed into a 4 x L binary matrix. L is either the WT length in the WT-specific model or 50 for the whole-library model.

##### Model evaluation

We performed 10-fold cross-validation to evaluate the binding models. We partitioned the dataset randomly into 10 equal-sized folds. Then, we trained and tested the model 10 times, each time using a different fold as the test set and the other nine folds combined as the training set. We used two measurements to gauge model performance: Pearson correlation and area under the ROC curve. Pearson correlation measures the linear agreement between two vectors, and is a common measure to evaluate intensity prediction. Area under the receiver operating curve (AUC) is a common measure to evaluate classification of positive and negative data points. We defined positive sequences as those having a binding intensity above 3.5 and negative sequences as those having a binding intensity of zero or less.

##### Parameters search

We used a hyper parameter search to optimize model performance. The first step of this search process was to randomly select a set of parameters from the parameter space defined for each of the models (Extended Data Tables 5 and 6), build, train and test the model. That step was repeated 10 times. Each random parameter combination was evaluated using 10-fold cross-validation as described above. From the 10 random parameters sets, the best performing set was selected based on average Pearson correlation over the 10 folds. The second step of the search was “fine tuning” of the chosen parameter set. In this step, we checked sets of parameters in the “surrounding” of the set that was selected during the first step. This process of parameters selection was done for each protein and for each of the model types separately.

#### WT-specific binding model

##### Dataset division

We first tackled the challenge of developing a model based on a WT and its variants of the same length. For this aim, we used a different subset of the data for each protein. The protein-specific subset contained only the sequences that have the same length as its WT binding site (MS2 – 19nt, Qβ – 20nt, PP7 – 25nt). Then, we split the subset again by the prefix of the sequence (C or GC). The rationale for the second split is the low correlation in binding intensities observed between C and GC positions (Extended Data Figure 4). This process is summarized in Figure 3A, main text.

##### Model description and optimization

Each WT-specific model is composed of 1-2 hidden layers with 10-40 nodes and one output layer with a single node (Figure 3A). Each protein and its sub-library have different parameters that were chosen specifically for it.

This optimization process was done as described under the Parameters search section above. The details of the parameters we examined are described in Extended Data Table 5.

**Extended Data Table 5.** Parameters search space for WT-specific model, The parameter space for each of the two steps of the hyper parameters search and the final models parameters. Unless noted otherwise, the range specified is of stride 1.

| Parameter space |  |  | Final parameters (protein, prefix) |  |  |  |  |  |
| --- | --- | --- | --- | --- | --- | --- | --- | --- |
| Parameter | Initial space | Surrounding space | MCP-C | MCP-GC | QCP-C | QCP-GC | PCP-C | PCP-GC |
| Nodes | 5-50 | $\pm 5$ | 38 | 32 | 39 | 39 | 10,10 | 10,10 |
| Layers | 1-3 | - | 1 | 1 | 1 | 1 | 2 | 2 |
| Activation function | linear, tanh, relu | - | tanh | tanh | tanh | tanh | tanh | tanh |
| Epochs | 20,30..100 | $\pm 15$ (strides of 5) | 70 | 70 | 70 | 70 | 70 | 70 |

In addition to the parameters in Extended Data Table, which are unique to each model, there are additional parameters that are common to all of them: learning rate 0.001 (default), batch size 8, optimizer ADAM, loss function MSE (mean squared error) and dropout of 0.2 for each hidden layer. The output layer consisted of one node with a linear activation function.

##### Evaluation

Overall, our WT-specific models achieved good prediction performance, i.e. average AUC of more or almost 0.6 and average Pearson correlation of more or almost 0.5, in 10-fold CV (Extended Data Figure 5). As explained before, the sub-library of each RBP was divided in to two sub-libraries based on its prefix. We trained a model specific for each of the two sub-libraries and tested it in 10-fold CV. We then choose the better performing model out of these two according to its average Pearson correlation in 10-fold CV, and used it in our downstream analysis. This resulted in using the C library for MCP and PCP, and the GC library for QCP.

#### Whole-library binding model

##### Padding sequences for whole-library models

We next developed a protein-specific binding model based on the whole library of RNA sequences and their responsiveness scores. Since the binding sites have different lengths, they need to be converted to have equal lengths for the learning process. All sequences were padded to the same length of 50nt. The binding sites were part of an RNA transcript. Hence, we upstream-padded them up to the transcription start site (TSS), with C or GC prefix according to their position. Downstream-padding of the sequences was done by their transcriptomic context up to a complete length of 50nt.

The upstream nucleotides are:

AATTGTGAGCGCTCACAATTATGATAGATTCAATTGGATTAATTAAAGAGGAGAAAGGTACCCATG

The downstream nucleotides are:

GTGAGCAAGGGCGAGGAGGATAACATGGCCATCATCAAGGAGTTCATGCGCTTCAAGGTGCACATGGAGGGCTCCGTGAACGGCCACGAGTTCGAGATCGAGGGCGAGGGCGAGGGCCGCCCCTACGAGGGCACCCAGACCGCCAAGCTGAAGGTGACCAAGGGTGGCCCCCTGCCCTTCGCCTGGGACATCCTGTCCCCTCAGTTCATGTACGGCTCCAAGGCCTACGTGAAGCACC

##### RNA secondary structure information

For the whole-library binding model, we augmented the one-hot encoded sequence information by RNA secondary structure information. We used RNAfold algorithm (Vienna package) to predict the structure of each sequence. The input to RNAfold is the binding site, and it outputs the predicted secondary structure in parenthesis notation, i.e. opening and closing parenthesis for base-pairs and a dot for unpaired nucleotide.

We converted this notation into an encoding of RNA structural contexts. This was done by a MATLAB script that encodes the RNA structure as a one-hot matrix with one bit set in each column for the corresponding structural context. For a binding site of length n, the n-long parenthesis annotation is transformed to a 5 x n binary matrix. The structural contexts we used were lower stem (LS), bulge (B), upper stem (US), loop (L), and no-hairpin (N). The one-hot encoded structure matrix not on the binding site was set to zero. The RNA structure matrix was concatenated to the sequence matrix, (Figure 4A, main text). In total, for a sequence of length L, this results in a binary matrix of size (4+5) x L.

##### Model description and optimization

The model is composed of one convolution layer, one hidden layer and an output layer (Figure 5A, main text).

The optimization of the model was done in the same manner as described above. Briefly, 10 random parameters sets were tested, and the best preforming one was chosen followed by fine tuning.

**Extended Data Table 6.**
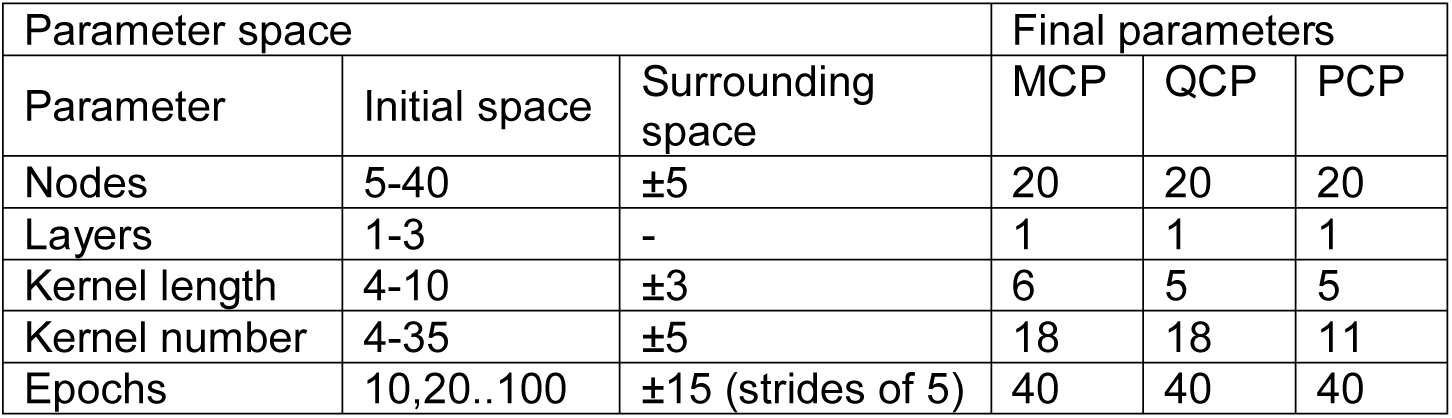
Parameters search for whole-library models. The parameter space for each of the two steps of the hyper parameters search and the final models parameters. Unless noted otherwise, the range specified is of stride 1.

| Parameter space |  |  | Final parameters |  |  |
| --- | --- | --- | --- | --- | --- |
| Parameter | Initial space | Surrounding space | MCP | QCP | PCP |
| Nodes | 5-40 | $\pm 5$ | 20 | 20 | 20 |
| Layers | 1-3 | - | 1 | 1 | 1 |
| Kernel length | 4-10 | $\pm 3$ | 6 | 5 | 5 |
| Kernel number | 4-35 | $\pm 5$ | 18 | 18 | 11 |
| Epochs | 10,20..100 | $\pm 15$ (strides of 5) | 40 | 40 | 40 |

In addition to the parameters in Extended Data Table 6, which are unique to each model, there are additional parameters that are common to all of them: learning rate 0.001, batch size 16, optimizer ADAM, loss function MSE (mean squared error), activation function for the convolution layer ‘relu’ and for the hidden layer ‘tanh’. The output layer consists of one node with a linear activation function.

##### Evaluation

The prediction performance achieved by the whole-library models are similar to the WT-specific ones, i.e. average AUC of more than 0.6 and average Pearson correlation of around 0.5 for each of the three proteins (Extended Data Figure 5). Although the WT-specific model achieved better results for two out of the three proteins, this whole-library model has the advantage that it can be applied to a binding site of any length, and not just that of the WT. This enables the prediction of binding of all three proteins to the same sequence set. To show case the contribution of RNA structure to our whole-library models, we trained and tested a model without the additional RNA structure information, and prediction performance degraded with the removal of RNA structural information (Extended Data Figure 6).

##### Generation of sequences for experimental validation

To test the predicted binding cassette generated according to our models’ predictions, we created one million synthetic binding sites. We generated one million random sequences that are in hamming distance of 3-7 from one of the WT binding sites. Overall, we randomly selected the one million out of 1.5 billion options. Because the number of possible variants rises as the length of the sequence, uniform selection of sequences will result in more variants of the long WT (PCP, 25-nt long) and less variants of the short WT (MCP, 19-nt). To overcome this bias, we divided the random selection into three parts; in each part we randomly selected 333,333 sequences from the variants of one WT. We computed the binding intensity of each of the proteins to the set of one million sequences using the whole-library models. Then, to experimentally validate model accuracy, we chose a sample out of the one million. We selected ten sequences that are single binders (i.e. bound by a single protein and not by the two others), and ten that are double binders (i.e. bound by two proteins and not by the third). All are in edit distance of at least 4 from one another and all were not included in the original experimental library.

## Supplementary items

5 figures

6 tables

5 movies

## Acknowledgements

The authors would like to acknowledge the Technion’s LS&E staff (Tal Katz-Ezov and Anastasia Diviatis) for help with sequencing.

## Author contributions

NK designed and carried out the oligo library and microscopy experiments and analysis for all constructs. ET wrote the code for the machine learning models and carried out the ML-based analysis. SG and OA assisted and guided the experiments. ZY assisted with the OL design and statistical analysis of results. RA and YO supervised the study. NK, RA, and YO, wrote the manuscript.

## Materials & Correspondence

