## Supplemental Figures for "Overcoming the design, build, test (DBT) bottleneck for synthesis of nonrepetitive protein-RNA binding cassettes for RNA applications"

### Supplementary figures

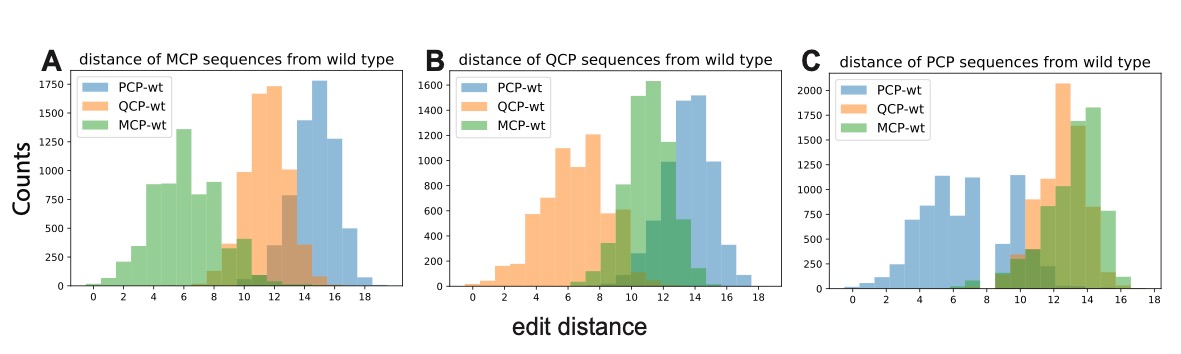

**Extended Data Figure 1.** Histograms of the edit distance of the sequences in the library of (A) MCP (B) QCP (C) PCP to the different wild types. The library contains sequences with high similarity to each of the wild types, with larger distances to the wild type of the other proteins.

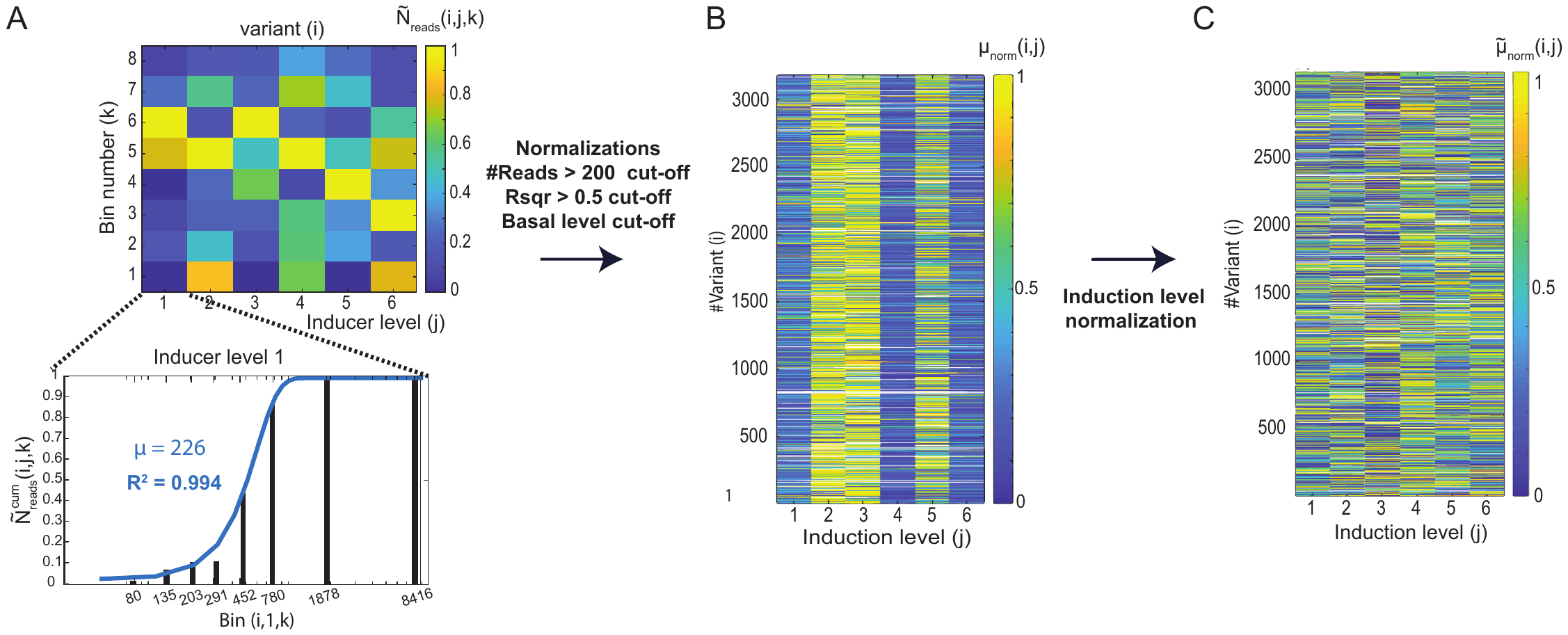

**Extended Data Figure 2.** Flowchart for the preliminary analysis conducted on the reads extracted from the oligo-library experiment. (A) (Top) a sample 6x8 matrix obtained for each variant. (Bottom) Collapsing the matrix to a vector of integrated mCherry level for every inducer value. (B) a sample list for PCP of unsorted non-renormalized 6-vectors displayed as heatmap. (C) Renormalized heatmap displaying unsorted PCP responsive variants.

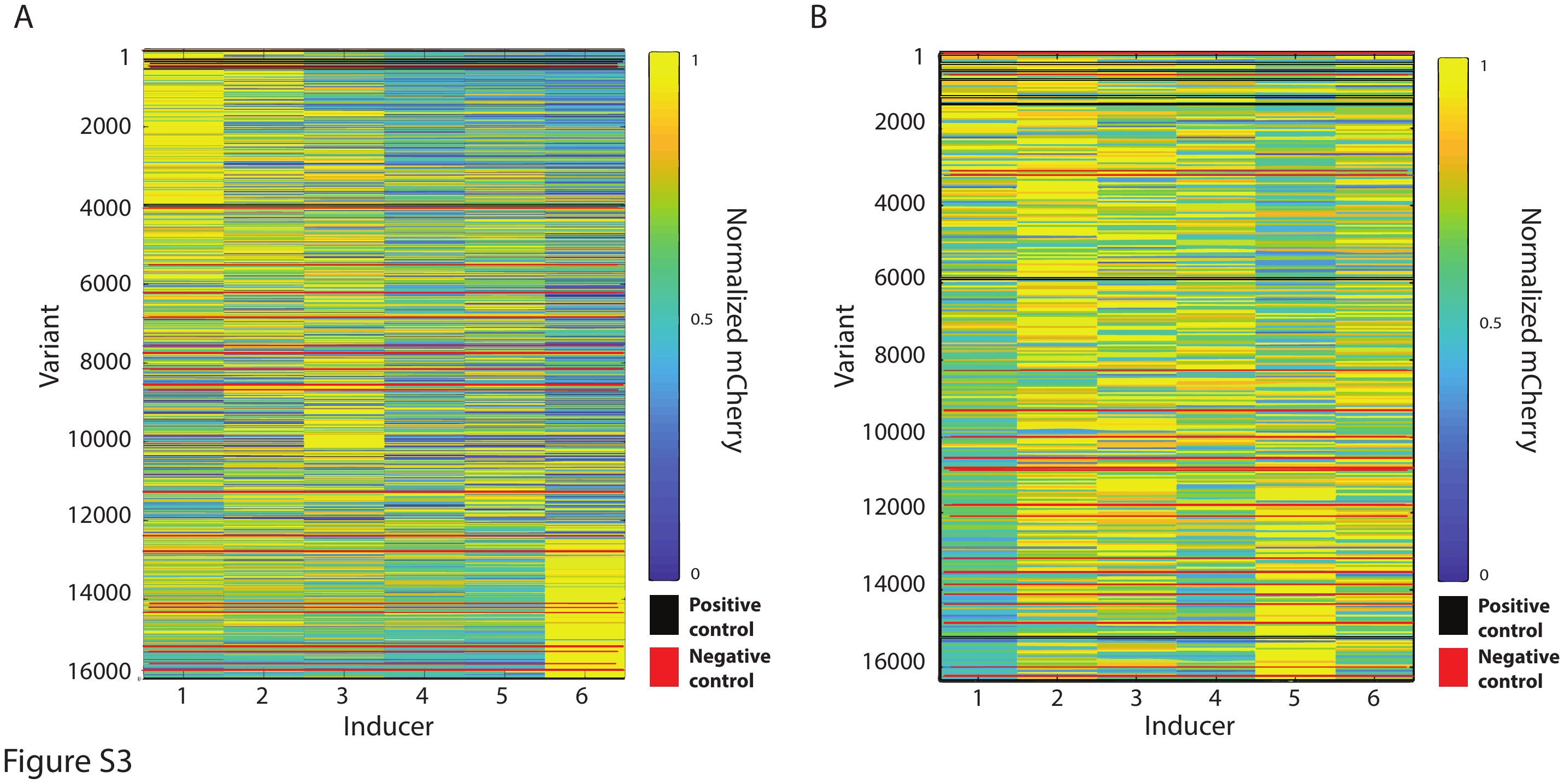

**Extended Data Figure 3.** Sorted heatmaps of MCP (left) and QCP (right) with the OL. Positive and negative control are depicted in black and red, respectively.

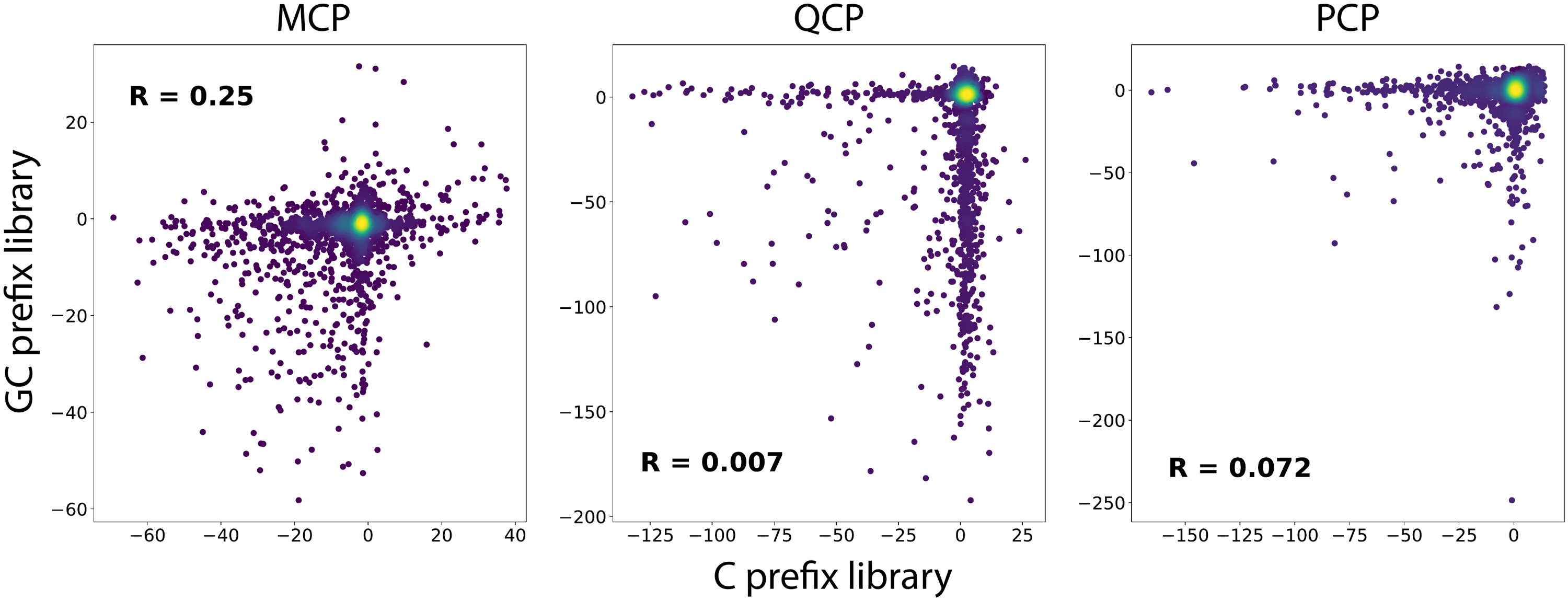

**Extended Data Figure 4**. Comparison of the binding scores between C and GC prefixes for the same binding sites for (A) MCP, (B) QCP, and (C) PCP. For all proteins, there is effectively no correlation between expression levels and the position of the variants within the ribosomal initiation region.

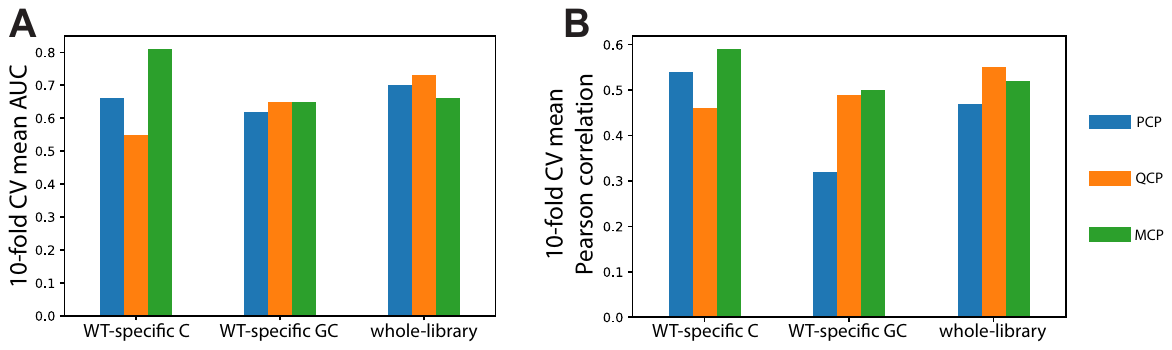

**Extended Data Figure 5.** The performances of the final WT-specific and whole library models. Performance accuracy is reported by (A) AUC, and (B) Pearson correlation.

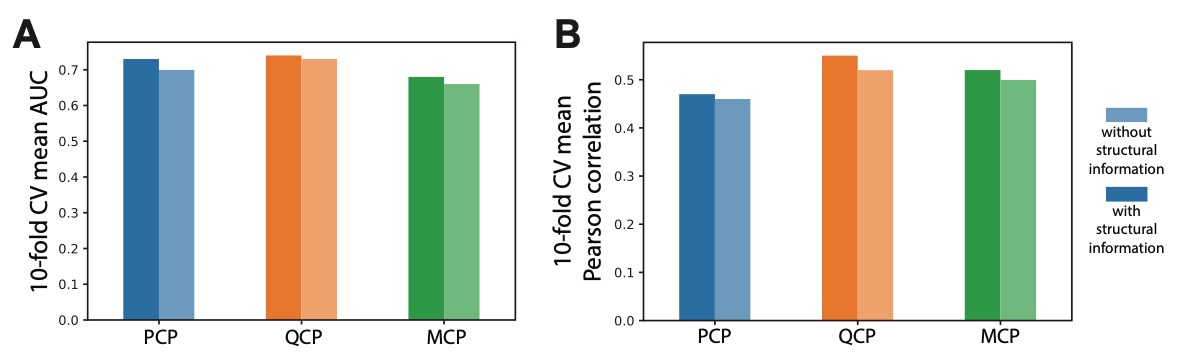

**Extended Data Figure 6** RNA structural information contribution to binding model prediction accuracy. Comparison of both (A) AUC and (B) Pearson correlation scores. The data shows that for all cases when the model was trained with structural information its performance improved.
